# The Nasal and Oropharyngeal Microbiomes of Healthy Livestock Workers

**DOI:** 10.1101/145029

**Authors:** Ashley E. Kates, Mark Dalman, James C. Torner, Tara C. Smith

## Abstract

Little information exists on the microbiomes of livestock workers. A cross-sectional, epidemiological study was conducted enrolling 59 participants (26 of which had livestock contact) in Iowa. Participants were enrolled in one of four ways: from an existing prospective cohort study (n=38), from the Iowa Department of Natural Resources Animal Feeding Operations database (n=17), through Iowa county fairs (n=3), and through snowball sampling (n=1). We collected two sets of swabs from the nares and oropharynx of each participant. The first set of swabs was used to assess the microbiome via 16s rRNA sequencing and the second was used to culture *S. aureus.* We observed livestock workers to have greater diversity in their microbiomes compared to those with no livestock contact. In the nares, there were 26 operational taxonomic units found to be different between livestock workers and non-livestock workers with the greatest difference seen with *Streptococcus* and *Proteobacteria.* In the oropharynx, livestock workers with swine exposure were more likely to carry several pathogenic organisms. The results of this study are the first to characterize the livestock worker nasal and oropharyngeal microbiomes.

## INTRODUCTION

The importance of microorganisms in maintaining human health has been recognized for many years. The composition of the microbiome is greatly influenced by ones environment [1]. It has been hypothesized the microbiome may protect those raised on farms from diseases such as asthma and atopy through animal-associated microbes and plant materials that stimulating the immune system and is known as the farm effect [2].

However, the farm effects ability to help protect against early disease is primarily seen in childhood. Adults working in close proximity to animals are at increased risk of respiratory conditions including chronic obstructive pulmonary disease (COPD), occupational asthma, and organic dust toxic syndrome. This is in part due to the inhalation of organic dust containing microorganisms [3, 4]. This is especially true for individuals working in enclosed animal houses as is common in swine and poultry production.

In order to better understand the relationship between the microbiome and livestock workers health, research is needed to characterize the microbiome of those with livestock contact. While research exists characterizing the air around livestock production facilities as well as the animals themselves, there is surprising limited information on the workers themselves. The aim of this study was to assess the microbial composition of the anterior nares and oropharynx of livestock workers compared to those without livestock contact using culture-independent techniques. To our knowledge, our study is the first to assess the microbiomes of the anterior nares and oropharynx of healthy livestock workers.

## METHODS

### Study Population

Participants were enrolled into a cross-sectional study between April 2015 and March 2016 in Eastern Iowa. Eligibility criteria were: 18 years of age, speak English, have not taken antibiotics or inhaled corticosteroids in the prior three months, not had the nasal influenza vaccine in the last month, no active infections of the upper respiratory tract, no hospitalized for greater than 24 hours in the last three months, and did not have HIV/AIDS. We also requested participants not eat, drink, or brush their teeth within one hour of sample collection.

Participants were enrolled in one of four ways. First, through a pre-existing cohort consisting of 95 families (177 participants over 18 years of age). One individual from each family was contacted by letter and then by phone call to schedule enrollment. If the original contact person for each family was either not interested or ineligible for participation, a letter was sent to the other members of the family unit until all eligible adults in the cohort were contacted. Only one individual from each family unit was eligible for participation. Participants enrolled from the pre-existing cohort were both livestock workers and non-livestock workers.

Livestock workers were also enrolled through the Iowa Department of Natural Resources (DNR) Animal Feeding Operations (AFO) database [5], Iowa county fairs, and snowball sampling. Operations were chosen from the DNR AFO database based on county (Johnson, Linn, Keokuk, Washington, and Louisa Counties) and mailed an invitation letter. One individual per AFO was eligible for enrollment. At the Iowa and Jones County fairs, a researcher passed out information on the study to livestock workers attending the fair. Participants could either take an information packet and contact the study team at a later date or could answer several eligibility questions and schedule an enrollment date while at the fair. Lastly, snowball sampling was used to recruit participants. Already enrolled livestock workers were asked to reach out to other livestock workers they knew (who did not live in their household and did not work on the same operation). The enrolled workers did not have to inform the study team how many packets were handed out or to whom. Interested potential participants then called the study team to set up enrollment. All study protocols were approved by the University of Iowa Institutional Review Board prior to enrollment.

### Sample Collection and Processing

Enrollment occurred in the participant’s home. After consenting, participants filled out questionnaires assessing demographic characteristics, medical history, and animal contact. Following the questionnaires, each participant provided swabs from their anterior nares and oropharynx. All samples were collected by a trained researcher and transported to the University of Iowa Center for Emerging Infectious Diseases (CEID) for processing. Samples were collected on sterile, dry, nylon flocked swabs (Copan Diagnostics, Murrieta, CA).

Bacterial DNA was isolated using the MO BIO PowerSoil DNA isolation kit (Mo BIO Laboratories Inc, Carlsbad, CA) adapted for swab use by removing the swab head and placing it in the tube during bead beating. Negative controls (kit reagents only) were used for every batch of extractions. Samples were sent for sequencing (including library preparation) to the University of Minnesota Genomics Center. 16s rRNA sequencing of the v1-v3 region was done on the Illumina MiSeq using 2x300 nt reads. Briefly, DNA was normalized to 5ng/μL for amplicon polymerase chain reaction (PCR) followed by a PCR clean-up step using AMPure XP beads to prepare for indexing. Index PCR was then done to attach the dual indices and sequencing adapters using the Nextera XT Index kit followed by another PCR clean-up step and library validation. Fluorometry was used for library quantification followed by normalization and pooling. The library was diluted to 4 nM and 5 μl of diluted DNA was used for pooling. The library was then denatured (using NaOH and heat) and diluted to prepare for sequencing on the MiSeq using the v3 chemistry. Primer sequences and PCR conditions can be found in the supplemental (Table S1).

### Statistical analysis

Sequences were assessed for quality using FastQC (Babraham Institute, Cambridge, UK) with poor quality reads filtered out (poor quality sequencing reads are defined as sequences with low base quality scores, short reads less than 200bp, reads with uncalled nucleotide bases, or any reads that could not assemble into paired reads). Reads were assembled using FLASh with the following parameters: minimum overlap = 30, maximum overlap = 150, and mismatch = 0.1 [6]. Adapters were removed from the merged file using Cutadapt [7]. USEARCH version 8.1.1861 and Python version 2.7.12 were used for chimera removal, operational taxonomic unit (OTU) binning, and taxonomy assignment at the genus level. The Ribosomal Database Project (RDP) classifier was used as the reference database. OTUs were grouped together based on 97% similarity. Any species level classification was done using BLAST+2.4.0 and the blastn function. Human-associated OTUs were also removed from the dataset using BLAST+2.4.0 and the blastn function. R version 3.3.1 was used for all statistical analyses and plot generation using the following packages: phyloseq [8], vegan [9], DESeq2 [10], and ampvis [11]. Alpha diversity was assessed using the Inverse Simpson diversity index [12] and beta diversity was assessed using the Bray-Curtis dissimilarity measure [13]. Principal coordinates analysis (PCoA) was used to visualize beta diversity. PERMANOVA, through the vegan package, was used to assess diversity differences between groups. PERMANOVA was chosen because it does not assume any distribution, unlike parametric tests [14]. The DESeq2 and ampvis packages were used to assess microbiota differences between groups. The DESeq2 package is only able to perform comparisons between two groups, as such animal contact was collapsed to swine versus all others when considering differentially abundant OTUs. Results were considered significant if the *P* was less than 0.05.

## RESULTS

### Participant demographics

Fifty-nine participants (26 livestock workers and 33 non-livestock workers) were enrolled (Figure 1). The average age of participants was 54.6 years (range: 28-85 years) and 41 (69.5%) were male. Livestock workers were significantly older than non-livestock workers (59.1 and 51.1 years respectively, *P*=0.027) and were predominantly male (92.3%) while males only made up 51.5% of the non-livestock workers (*P* = 0.0007) (Table 1). Those without livestock contact were more likely to brush their teeth daily (*P* <0.001), use liquid hand soaps (*P* <0.001), and more likely to use a gym (*P*=0.011) compared to those with livestock contact. (Table 2). There were no other significant differences between those with and without livestock contact.

**Figure 1:**
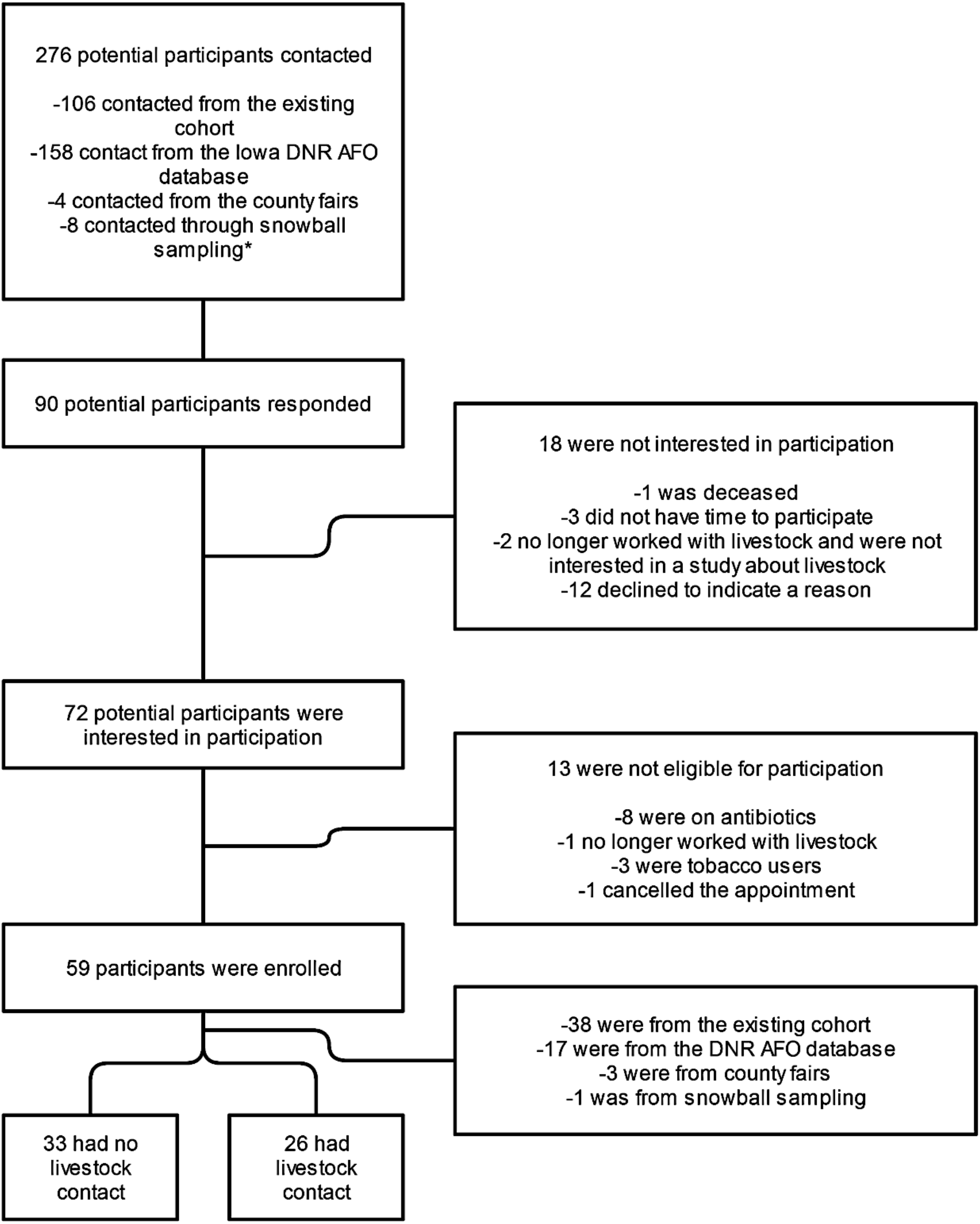
Flow diagram of participant enrollment.

**Table 1:**
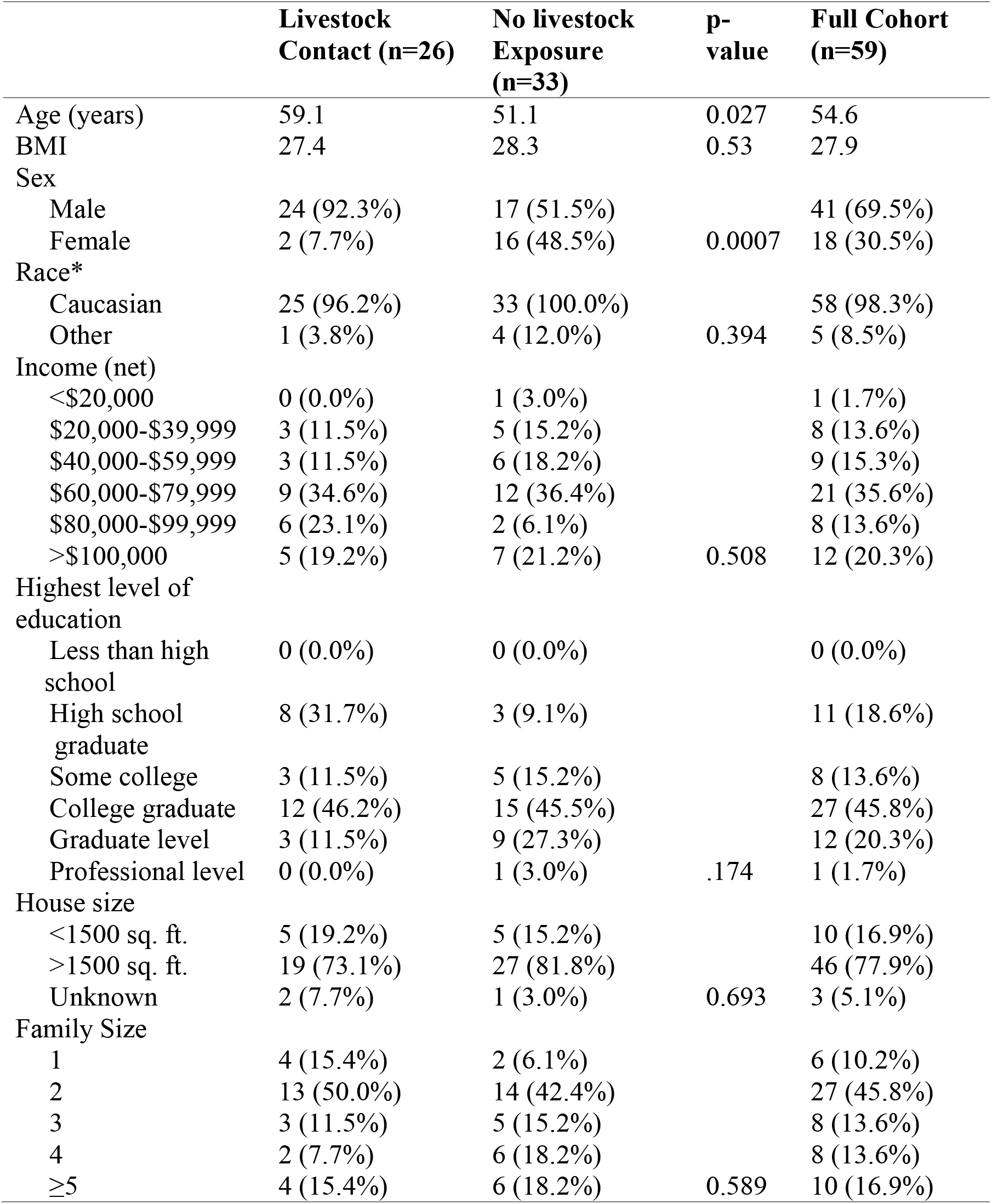
Participant demographics

**Table 2:** Health and Hygiene characteristics of participants

|  | **Livestock**<br>**Contact (n=26)** | **No livestock**<br>**Exposure (n=33)** | **p-**<br>**value** | **Full Cohort**<br>**(n=59\*)** |
| --- | --- | --- | --- | --- |
| Asthma |  |  |  |  |
| Yes | 0 (0.0%) | 3 (9.1%) |  | 3 (5.1%) |
| No | 25 (100.0%) | 30 (90.0%) | 0.25 | 55 (93.2%) |
| COPD |  |  |  |  |
| Yes | 1 (4.2%) | 1 (3.1%) |  | 2 (3.5%) |
| No | 23 (95.8%) | 32 (96.9%) | 0.429 | 55 (96.5%) |
| Heart Disease |  |  |  |  |
| Yes | 2 (8.0%) | 6 (18.2%) |  | 8 (13.8%) |
| No | 23 (.92%) | 27 (81.8%) | 0.445 | 50 (86.2%) |
| Diabetes |  |  |  |  |
| Yes | 1 (4.0%) | 1 (3.0%) |  | 2 (3.4%) |
| No | 24 (96.0%) | 32 (97.0%) | 1.0 | 56 (96.6%) |
| Cancer |  |  |  |  |
| Yes | 1 (4.0%) | 3 (10.0%) |  | 5 (9.1%) |
| No | 24 (96.0%) | 26 (86.7%) |  | 49 (89.1%) |
| Don't know | 0 (0.0%) | 1 (3.3%) | 0.158 | 1 (1.8%) |
| Past Cigarette Use |  |  |  |  |
| Yes | 2 (8.0%) | 9 (27.3%) |  | 11 (19.0%) |
| No | 23 (92.0%) | 24 (72.7%) | 0.09 | 47 (81.0%) |
| Past Cigar Use |  |  |  |  |
| Yes | 0 (0.0%) | 4 (12.1%) |  | 4 (6.8%) |
| No | 26 (100.0%) | 29 (87.9%) | 0.123 | 55 (93.2%) |
| Past Chew User |  |  |  |  |
| Yes | 2 (7.7%) | 3 (9.1%) |  | 5 (8.5%) |
| No | 24 (92.3%) | 30 (90.9%) | 0.848 | 54 (91.5%) |
| Dentures |  |  |  |  |
| Yes | 0 (0.0%) | 2 (6.3%) |  | 2 (3.4%) |
| No | 26 (100.0%) | 30 (93.8%) | 0.497 | 56 (96.6%) |
| Tooth Brushing<br>Frequency |  |  |  |  |
| Every morning | 9 (34.6%) | 26 (78.8%) |  | 35 (59.3%) |
| Every evening | 17 (65.4%) | 15 (45.5%) |  | 32 (54.2%) |
| Most mornings | 2 (7.7%) | 1 (3.0%) |  | 3 (5.1%) |
| Most evenings | 4 (15.4%) | 4 (12.1%) |  | 8 (13.6%) |
| Some mornings | 6 (23.1%) | 1 (3.0%) |  | 7 (11.9%) |
| Some evenings | 2 (7.7%) | 1 (3.0%) |  | 3 (5.1%) |
| No mornings | 0 (0.0%) | 0 (0.0%) |  | 0 (0.0%) |
| No evenings | 0 (0.0%) | 0 (0.0%) | <0.001 | 0 (0.0%) |
| Probiotic usage |  |  |  |  |
| Yes | 3 (11.5%) | 4 (12.1%) |  | 7 (12.1%) |
| No | 23 (88.5%) | 29 (87.9%) | 1.0 | 51 (87.9%) |

Table 2 continued: Health and Hygiene characteristics of participants
|  | **Livestock**<br>**Contact (n=26)** | **No livestock**<br>**Exposure (n=33)** | **p-**<br>**value** | **Full Cohort**<br>**(n=59\*)** |
| --- | --- | --- | --- | --- |
| Type of Hand Soap |  |  |  |  |
| Non-antibacterial, bar | 11 (42.3%) | 12 (36.4%) |  | 23 (40.0%) |
| Non-antibacterial, liquid | 11 (42.3%) | 17 (51.5%) |  | 28 (47.5%) |
| Antibacterial, bar | 11 (42.3%) | 8 (24.2%) |  | 19 (32.2%) |
| Antibacterial, liquid | 11 (42.3%) | 19 (57.6%) |  | 30 (50.8%) |
| Other | 1 (3.8%) | 0 (0.0%) | 0.001 | 1 (1.7%) |
| Visited a Correctional Facility |  |  |  |  |
| Yes | 1 (%) | 1 (%) |  | 2 (3.4%) |
| No | 25 (%) | 32 (%) | 1.0 | 56 (96.6%) |
| Outpatient surgery in last 3 months? |  |  |  |  |
| Yes | 0 (0.0%) | 1 (%) |  | 1 (1.7%) |
| No | 26 (100.0%) | 31 (%) | 1.0 | 57 (98.3%) |
| Visited a hospital or long-term care facility? |  |  |  |  |
| Yes | 14 (%) | 10 (%) |  | 24 (41.4%) |
| No | 12 (%) | 22 (%) | 0.142 | 34 (58.6%) |
| Work/ volunteer in a healthcare facility? |  |  |  |  |
| Yes | 1 (%) | 7 (%) |  | 8 (14.0%) |
| No | 25 (%) | 24 (%) | 0.59 | 49 (86.0%) |

Twenty-six participants had current exposure to livestock (Table 3).The majority of participants worked with swine (n=18). Several participants currently worked with more than one type of animal with seven participants working with two animal types, two working with three animal types, and one participant working with five animal types (swine, poultry, cattle, sheep, goats, and horses). The most frequent combination of animal types was swine and cattle (n=4).

**Table 3:** Livestock contact (n=26)

| **Animal** | **N (%)** | **Ave. Number**<br>**Animals (range)** | **Ave. Days per**<br>**week (range)** | **Ave. Hours per**<br>**day (range)** |
| --- | --- | --- | --- | --- |
| Swine | 18 (69.2%) | 3,024 (8-10,000) | 6 d (2-7) | 2.5 h (0.5-10) |
| Cattle | 12 (46.2%) | 191 (4-850) | 6.4 d (1-7) | 1.6 h (0.5-3) |
| Poultry | 4 (15.4%) | 1,644 (20-6,500) | 7 d | 1.0 h (0.25-2) |
| Other |  |  |  |  |
| Sheep | 4 (15.4%) | 28 (10-50) | 6.5 d (6-7) | 1.4 h (0.25-3) |
| Horses | 2 (7.7%) | 6.5 (1-12) | 6.5 d (6-7) | 5.5 h (3-8) |
| Goats | 1 (3.8%) | 10 | 7 d | 2 h |

### Microbiota analysis

The Inverse Simpson diversity index (Figure 2a) was greater for those with livestock contact compared to those without livestock contact in the nasal samples (p > 0.001); however, there was no difference in the oropharyngeal samples (p = 0.542). The ordination plot of the Bray-Curtis distances for all samples is shown in Figure 2b. The samples cluster by sample type (*P* = 0.001) and livestock exposure (*P* = 0.038, *P* for the interaction between sample type and livestock exposure = 0.035). Because samples cluster by both livestock exposure and sample type, the nasal (Figure 2c) and oropharyngeal samples (Figure 2d) were assessed separately. A significant difference remained in the nasal samples (*P* = 0.002), but not the oropharyngeal samples (P= 0.559). There were no differences in the diversity of the microbiomes based on any participant characteristics (data not shown).

**Figure 2:**
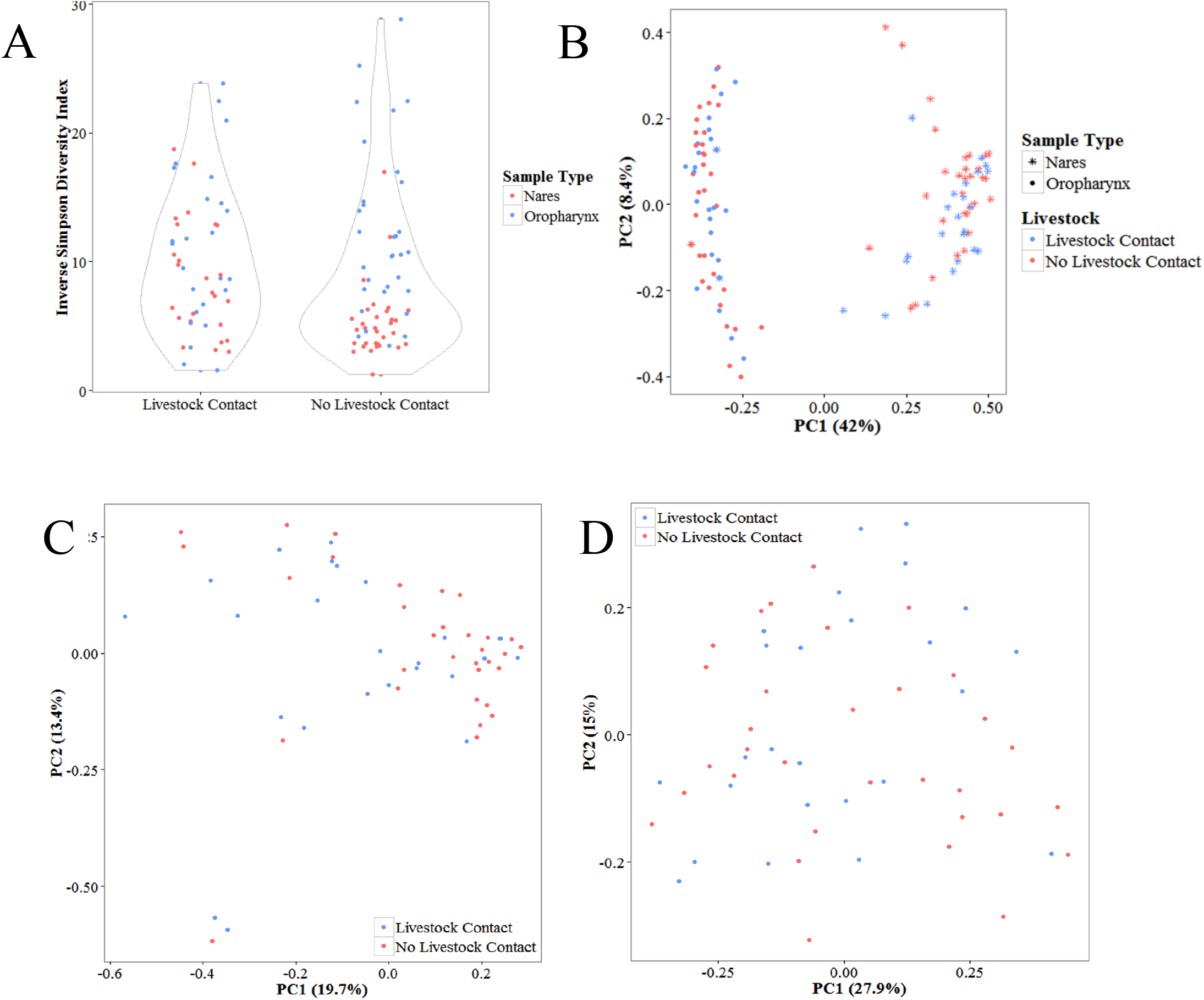
Diversity analysis by livestock contact. a) alpha diversity of both nasal and oropharyngeal samples by livestock exposure. b) PCoA of the Bray-Curtis dissimilarity matrix all samples by livestock exposure. c) PCoA of the nasal samples. d) PCoA of the oropharyngeal samples. PC1 and PC2 = principle coordinates 1 and 2 respectively.

There was no difference in alpha diversity by animal type (cattle, poultry, swine, more than one animal type) in either the nares (*P* = 0.762) or oropharynx (*P* = 0.941). In the nares, there was a difference by animal types (*P* = 0.009); however, there are no differences in the oropharynx (*P* = 0.297).

Actinobacteria and Firmicutes were the most prevalent phyla in both the livestock workers and non-livestock workers. Bacteroidetes were more abundant in the livestock workers. The barplot and boxplot of the most abundant OTUs can be found in the supplemental (Figures S3, S4). A total of 26 OTUs were differentially represented between the livestock workers and non-livestock workers, 25 of which were significantly more abundant in those with livestock contact. Only two OTUs belonging to the *Streptophyta* genus were more abundant in the non-livestock workers (Figure 3).

**Figure 3:**
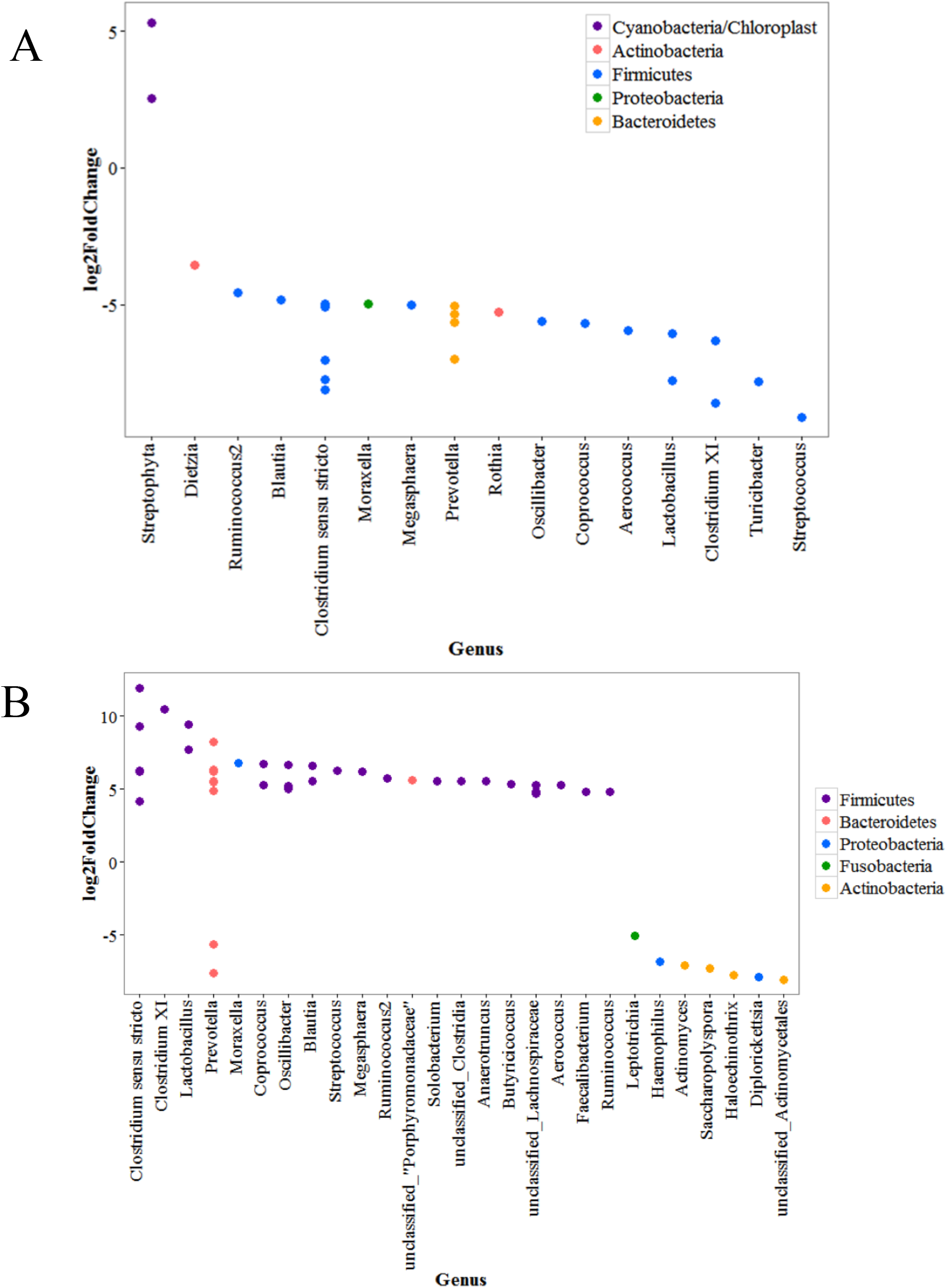
Log 2-fold Change of the significantly differentially abundant OTUs (Benjamini-Hochberg correction applied). Points represent OTUs with phyla represented by color. Negative values represent OTUs significantly more abundant in livestock workers and positive values represent OTUs significantly more abundant in non-livestock workers. a) Differentially abundant OTUs in the nares and b) differentially abundant OTUs in the nares by animal contact (swine vs. all others).

Unlike the nasal microbiome, there is a great deal of similarity between those with and without livestock contact in the oropharynx. There were no OTUs significantly differentially abundant between the livestock workers and those without livestock contact. The *Streptococcus* genera was the most prevalent genus observed in the oropharynx followed by *Provetella* and *Heamophilus* genera.

When stratifying by animal type in the nares, *Corynebacterium* and *Staphylococcus* were the most abundant genera with members of the Firmicutes phylum being the most abundant. When comparing swine workers to those with any other animal contact, one OTU was significantly more abundant in the swine workers, *Clostridium sensu stricto* (2-fold change: 8.58, *P* < 0.001). In the oropharynx there were nine OTUs significantly more abundant in the swine workers compared to those with all other animal types and two *Lactobacillus* OTUs with increased abundance in those with no swine contact (Figure 4).

**Figure 4:**
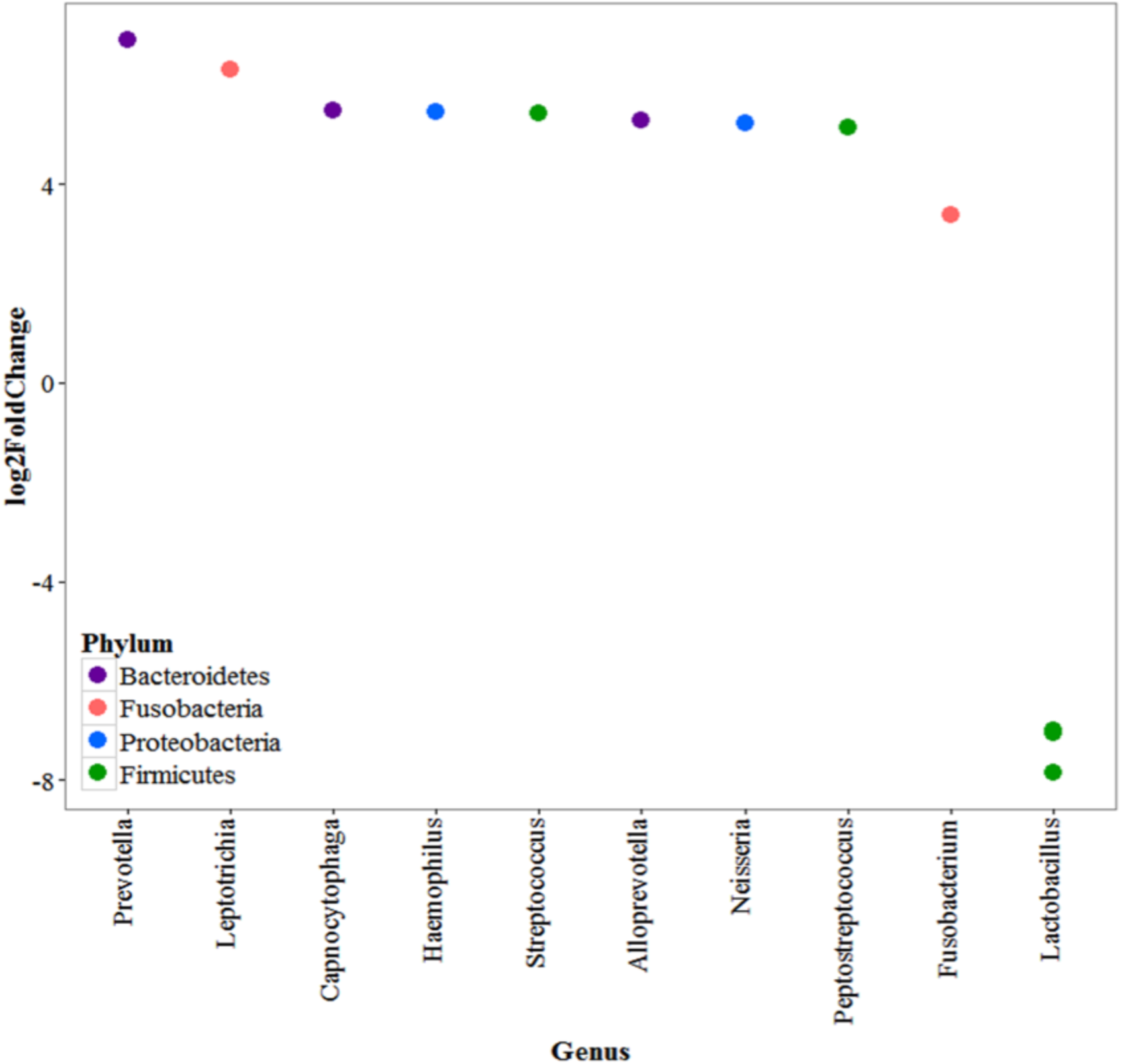
Log 2-fold Change of the significantly differentially abundant OTUs in the oropharynx by animal contact (swine vs. all others). (Benjamini-Hochberg correction applied). Points represent OTUs with phyla represented by color. Negative values represent OTUs significantly more abundant in livestock workers and positive values represent OTUs significantly more abundant in non-livestock workers

## DISCUSSION

Very little is known about the healthy livestock worker nasal and oropharyngeal microbiomes. The majority of studies assessing the microbial communities related to livestock work have either been done in animals [15, 16] or have studied the aerosolization of microorganisms in and around livestock facilities [3, 4, 17]. Here we have described the nasal and oropharyngeal microbiomes of 26 livestock workers and 33 non-livestock workers in Iowa.

The population was comprised of primarily older (mean age of 54.6 years), Caucasian (98.3%) males (69.5%). Those with livestock contact were significantly older than those without livestock contact (59.1 years compared to 51.1 years) as well as more likely to be male (92.3% male compared to 51.5% male). This represents the average farmer worker in the United States where a majority of farm workers are males [18]. In the majority of Iowa counties, including Keokuk County, less than 10% of farm workers are female. Additionally, we observed no microbiota differences between males and females (data now shown). Furthermore, as of 2012 the average age of principal farmworkers was 58.3 years with 61% being between 35 and 64 years nationwide [18].

The importance of livestock contact on the human microbiome has been recognized in relation to respiratory diseases. It has been suggested that the farm effect is protective against asthma. This is particularly true for children where it has been shown early life exposure to microbes and microbial components prime the immune system by the upregulation of T-helper 1 cells and the downregulation of T-helper 2 cells reducing the risk of atopy [19]. Studies have shown having a parent in a farming occupation – particularly ones with livestock exposure – is significantly associated with lower rates of allergen disorders and allergy attacks and there is a dose response relationship with less atopy in children with parents who are full-time farmers [20, 21]. It is thought the high-diversity of microorganism – likely inhaled – outcompete the harmful bacteria that may promote asthma [2]. In adults farmer’s asthma is low (around 4%) as is atopy (14%); however, unlike in children, asthma rates are higher among those who work with livestock, particularly swine and cattle [22]. Studies have also shown asthma to be more common in farmers without atopy than those with atopy and individuals with more than one type of animal exposure were at increased risk of non-atopic asthma [22].

Livestock workers had significantly more diverse nasal microbiomes compared to non-livestock workers likely due to inhalation. Livestock workers are exposed to high levels of inhalable dust which contains microorganisms [3, 4]. The *Ruminococcaceae* family and *Lactobacillus* which were both found to be significantly more abundant in the nares of those participants with livestock contact than those lacking this exposure, have been identified in inhalable dust [23]. *Moraxella* – a human commensal also known to cause respiratory tract infections [24] – is a bacterial air contaminant in livestock houses [25]. Others have found organisms belonging to the *Aerococcaceae* family, *Dietzia,* and *Prevotella* in air surrounding livestock [17]. OTUs belonging to all of these genera were significantly more abundant in the nares of those with livestock contact in our population leading to the conclusion these organism may be being inhaled.

We identified several potential pathogens as more abundant in livestock workers’ nares and oropharynx. One of the organisms found to be significantly more abundant in the livestock worker microbiome was *Dietzia,* a gram positive genus known to be an opportunistic pathogen and able to colonize skin and formerly classified as *Rhodococcus maris* [26]. It is unsurprising that this genus is also able to colonize the anterior nares, as they are anatomically similar to the skin [27]. *Dietzia* is predominantly a zoonotic pathogen, but has been identified in invasive human infections as well [28–30]. Due to its similarity to *Rhodococcus* spp., it is often mistakenly identified as a contaminant [26, 31]. *Dietzia* was found to be roughly seven times more abundant in livestock workers compared to those with no livestock contact (2-fold change of -3.55) in our population. While *Dietzia* was found in the negative controls, it was found in few samples and likely was not a large enough contaminant to account for the large difference between the groups. *Dietzia* infection has been thought to be potentially related to prior livestock exposure in case reports [32] and has been identified in the air of poultry (duck) barns [33]. Due to its high prevalence in livestock workers, it may be a potential cause of difficult-to-diagnose infections in people with livestock contact, especially in the immunocompromised [34]; however, little information on *Dietzia* as an opportunistic pathogen exists. Other potential pathogens found in higher abundance in livestock workers were *Prevotella* [35–37], *Streptococcus* [38–40], *Moraxella* [41, 42], *Rothia* [43], and *Oscillibacter* [44].

*Prevotella* spp., particularly *P. ruminicola,* are difficult to culture microorganism prevalent in the gastrointestinal tracts of all livestock animals in addition to ruminants. It has been demonstrated *P. ruminocola* has the ability to transfer tetracycline resistance to other members of the Bacteroidetes phylum, particularly to other *Prevotella* species, in the host and horizontal transfer of the *tetQ* gene among *Prevotella* spp. is common in the human and ruminant intestines as well as the human oral cavity [45]. While it was not significantly enhanced in the livestock worker microbiome, *P. ruminocola* was present as were many oral-associated *Prevotella* species. *Prevotella* spp. are frequent causes of odontogenic infections associated with gram-negative, anaerobic bacteria [46, 47]. These organisms are also known to cause infections of the respiratory system, head, and neck [47]. This is of interest as tetracycline is still commonly used in agriculture as well as a treatment for periodontal disease [48, 49] and *Prevotella* spp. were very common in the nares and oropharynx in our population and significantly more abundant in the oropharynx of swine workers.

As it is likely these organisms are being inhaled while working around livestock, it is possible their presence is contamination and not true colonization. While there is little research surrounding contamination vs. colonization, several studies have been done with regard to livestock worker colonization with *S. aureus* and have found many livestock workers drop *S. aureus* carriage within 24 hours [50]. On average it had been roughly 30 hours since swine workers had their last contact with swine, 24 hours since cattle workers had their last contact with cattle, and 1.5 hours since poultry workers had their last contact with poultry at the time of swabbing. It is possible some of the organisms observed in the nasal microbiome were due to contamination from recently being around their livestock, especially in those with poultry contact. As many of the swine and cattle workers were close to 24 hours since their last contact with animals, it is difficult to determine if the presence of these organisms is true colonization or temporary contamination without further longitudinal research.

We observed three participant behaviors to be significantly different between those with and without livestock contact: type of soap used, gym usage, and the frequency of tooth brushing. However, none of these behaviors were significantly associated with alterations in either the nasal or oropharyngeal microbiomes. The most surprising of these was that frequency of tooth brushing, which was less frequent in the livestock workers, but was not associated with any differences in oral microbiota. One explanation for this is frequency of tooth brushing may not be an adequate marker of oral hygiene. While we chose to assess oral health through a single question (frequency of tooth brushing) in this pilot study as the enrollment visit was already long and required participants to fill out up to three surveys, in future studies directed towards assessing oral health and the livestock worker microbiome, this will not be sufficient. A better marker for oral hygiene may have been to assess the number of dental carries, gingivitis, gum disease, and/or halitosis. In the future, it would be better to assess oral hygiene using a standardized survey, such as the NHANES Oral Health Survey [51].

Our study is the first we are aware of to assess the microbiome of livestock workers using next-generation sequencing technology and great deal of additional research is needed. More research is needed to better understand the relation of the livestock worker respiratory microbiomes and diseases such as asthma. Longitudinal studies need to be done to first characterize the livestock workers over time and at different stages of life. Animal-based studies are needed to more definitively assess the relationship between the core microbes of the livestock worker airways and their impact on asthma. Animal models are necessary for this research to be able to determine if the microbes encountered during early childhood exposure to farm-life may be able to prevent asthma.

## Acknowledgments/funding

The authors would like to thank Dr. Patrick Breheny for his invaluable guidance as well as the University of Minnesota Genomics Center for conducting the sequencing. This publication was supported in part by Grant Number 5 U54 OH007548-11 from CDC-NIOSH. Its contents are solely the responsibility of the authors and do not necessarily represent the official views of the CDC, NIOSH, or the Great Plains Center for Agriculture Health.

